# Genetic basis of offspring number and body weight variation in *Drosophila melanogaster*

**DOI:** 10.1101/2020.07.14.203406

**Authors:** Jamilla Akhund-Zade, Shraddha Lall, Erika Gajda, Denise Yoon, Benjamin de Bivort

**Author notes:** equal contribution (Shraddha Lall, Erika Gajda).

## Abstract

*Drosophila melanogaster* egg production, a proxy for fecundity, is an extensively studied life-history trait with a strong genetic basis. As eggs develop into larvae and adults, space and resource constraints can put pressure on the developing offspring, leading to a decrease in viability, body size, and lifespan. Our goal was to map the genetic basis of offspring number and weight under the restriction of a standard laboratory vial. We screened 143 lines from the *Drosophila* Genetic Reference Panel for offspring numbers and weights to create an ‘offspring index’ that captured the number vs. weight trade-off. We found 30 associated variants in 18 genes. Validation of *hid*, *Sox21b*, *CG8312*, and *mub* candidate genes using gene disruption mutants demonstrated a role in adult stage viability, while mutations in *Ih* and *Rbp* increased offspring number and increased weight, respectively. The polygenic basis of offspring number and weight, with many variants of small effect, as well as the involvement of genes with varied functional roles, support the notion of Fisher’s “infinitesimal model” for this life-history trait.

## Introduction

Life-history traits, such as fecundity, lifespan, and body size, are major contributors to fitness. In *Drosophila melanogaster,* the genetics, plasticity, and evolution of life-history traits have been extensively studied^1^. *Drosophila* fecundity, measured through egg-laying behavior, was previously shown to have a strong genetic component that differs between young and old flies^2–5^, and is also influenced by temperature and nutrition^6,7^. Fecundity interacts with a number of physiological processes. In support of an energy allocation model of life-history^8^, fecundity has been shown to trade-off with longevity^5,9^, indicating that investment into the next generation can come at a cost to somatic maintenance via a transfer of energy reserves. A genome-wide association study revealed that age-specific fecundity is associated with variants present across a large set of candidate genes, enriched for genes involved in development, morphogenesis, neural function, and cell signaling^4^. Connecting fecundity with neural function, quantitative trait locus and deficiency mapping revealed that expression of a *Drip* aquaporin in corazonin neurons was positively correlated with fecundity by modulating the neurohormone balance between corazonin and dopamine^10^.

While the vast majority of *Drosophila* fecundity studies have used egg production as a measure of fecundity, the number of eggs laid may not translate perfectly to viable offspring due to potential mortality at the larval stages. Under both natural and laboratory conditions, larvae must contend with a finite space and resource limitations given the constraints of the rotting food substrate^11,12^ or culture media, as well as competition between larvae. Increased larval density decreases egg-to-adult viability^13–15^, body size^14,16–19^ and longevity20, while increasing development time^14,15,17,19^ and lowering starvation resistance^16^. Highly fecund flies that lay a large number of eggs may end up negatively affecting their offspring due to the increased larval density. On the other end of the spectrum, flies producing fewer eggs may have large offspring capable of weathering stress^21,22^, but fewer offspring result in a decreased competitive ability with more fecund individuals. Moreover, larvae need a critical number to engage in cooperative food-burrowing^23^, with cooperative food-burrowing more likely to occur in conspecifics of high relatedness^24^. Taken together, these pressures mean that finite space and resource limitation likely impose a trade-off between the number of offspring and their body phenotypes.

Given that these tradeoffs come into play after egg-laying, we were interested in whether there was a genetic basis for adult offspring number and their quality under resource limitation. We used the standard laboratory food vial to impose both a space and food limitation on the developing offspring. As a measure of quality, we measured the wet weight of recently eclosed offspring; increased body weight is correlated with increased starvation resistance^22^, increased nutrient stores, and increased immunity^25^, which indicate an investment in somatic maintenance. We scored lines from the *Drosophila* Genetic Reference Panel (DGRP)^26^ for numbers of adult offspring and their weight. In a genome-wide association analysis, we found candidate genes with variants significantly associated with a combined metric of offspring number and weight. Mutation of these genes, in most cases, caused lethality or impaired survival at the adult stage, but in other cases shifted the balance between offspring weight and number.

## Results

### Genome-wide associations of an offspring life-history index

We collected four fecundity and body weight phenotypes from 143 DGRP lines: total number of female progeny, total number of male progeny, and their respective mean weights (in mg). We found inter-line differences for all four phenotypes measured (Fig. 1a), as well as strong correlations between the phenotypes (Fig. 1b). The estimated broad-sense heritability of mean female weight (0.64, 95% highest posterior density interval (HPDI): 0.55 - 0.72) was lower than for mean male weight (0.73, 95% HPDI: 0.65 - 0.80). Both weight phenotype heritabilities were higher than previously estimated heritabilities for body weight^22,27^. The heritabilities of the total number of female progeny (0.47, 95% HPDI: 0.37 - 0.57) and male progeny (0.48, 95% HPDI: 0.39 - 0.58) were higher than the heritabilities previously estimated on number of eggs laid^4^ (Supp. Table 1).

**Figure 1.**
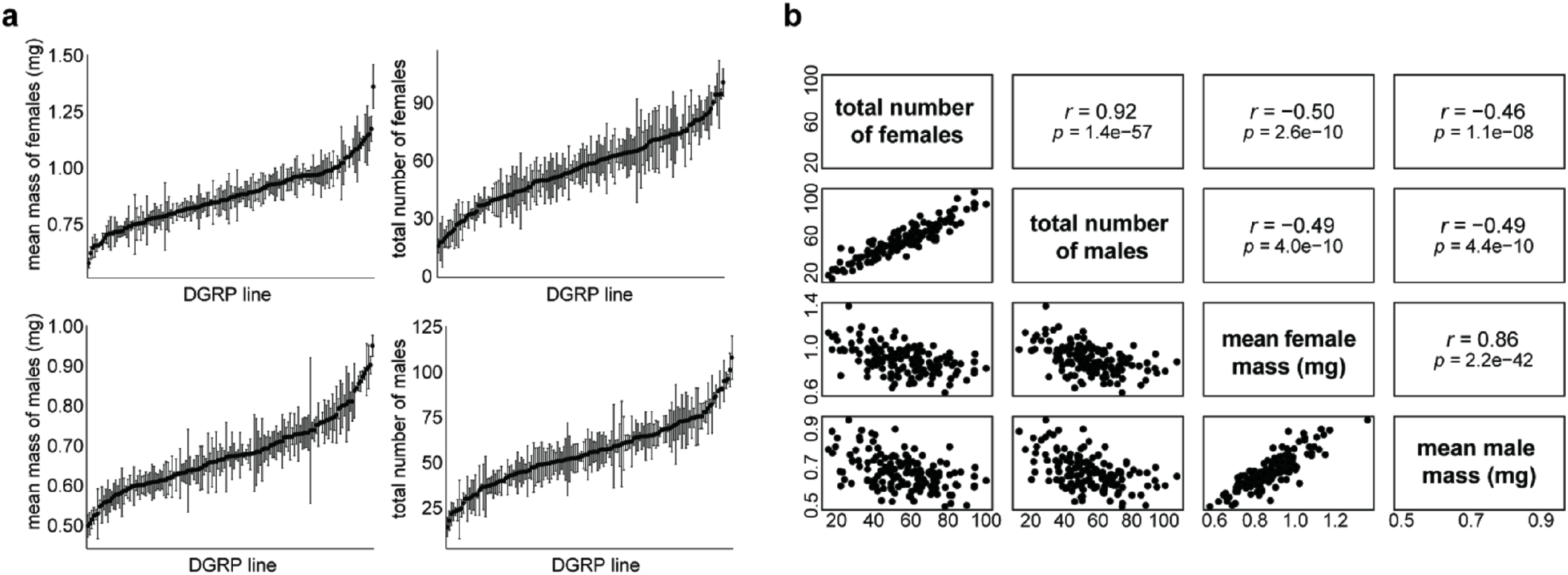
DGRP lines show variation in offspring number and weight. **a)** Plot of phenotypes measured (+/− 1 s.e.) with the DGRP lines sorted by the mean value for each phenotype (3 replicates/line). **b)** Correlation matrix of the phenotypes measured. Points are DGRP genotypes.

As we predicted based on tradeoffs imposed by resource limitation, we found that the number of offspring was negatively correlated with the offspring weight (Fig. 1b). The strong correlations between the offspring number and weight phenotypes allowed us to use principal component analysis to combine the four measurements into a single metric, which we termed the offspring index. The first principal component explained 71% of the variance in the data and is negatively loaded for offspring number and positively loaded for offspring weight (Supp. Table 2). A negative index value indicated many low-weight offspring, a positive index value indicated few high-weight offspring, and an index value close to zero indicated a balance between offspring number and weight.

We used this offspring index for a genome-wide association study. Thirty variants were associated with the offspring index using a threshold of *p* < 10^-5^ (Table 1). The third chromosome had 12 significant variants, while 16 were located on the second chromosome and only 2 on the X chromosome (*χ^2^* test, *p* = 0.32). Most of the variants (23/30) were within 1,000bp of a gene; roughly half of these (14/23) were located in introns. Five of the associated variants were present in genes previously associated with fecundity^4^, which is more overlap than expected given a random set of candidate genes (Supp. Fig. 1a). Among the candidate genes, we did not find significant enrichment for particular biological processes or molecular functions using PANTHER’s Overrepresentation Test with the GO-Slim annotation sets^28^. The QQ plot showed no systematic bias and a slight enrichment for *p* < 10^-5^ (Supp. Fig. 1b). The linkage disequilibrium heat map revealed no long-distance linkage between variants (Supp. Fig. 1c).

**Table 1.**
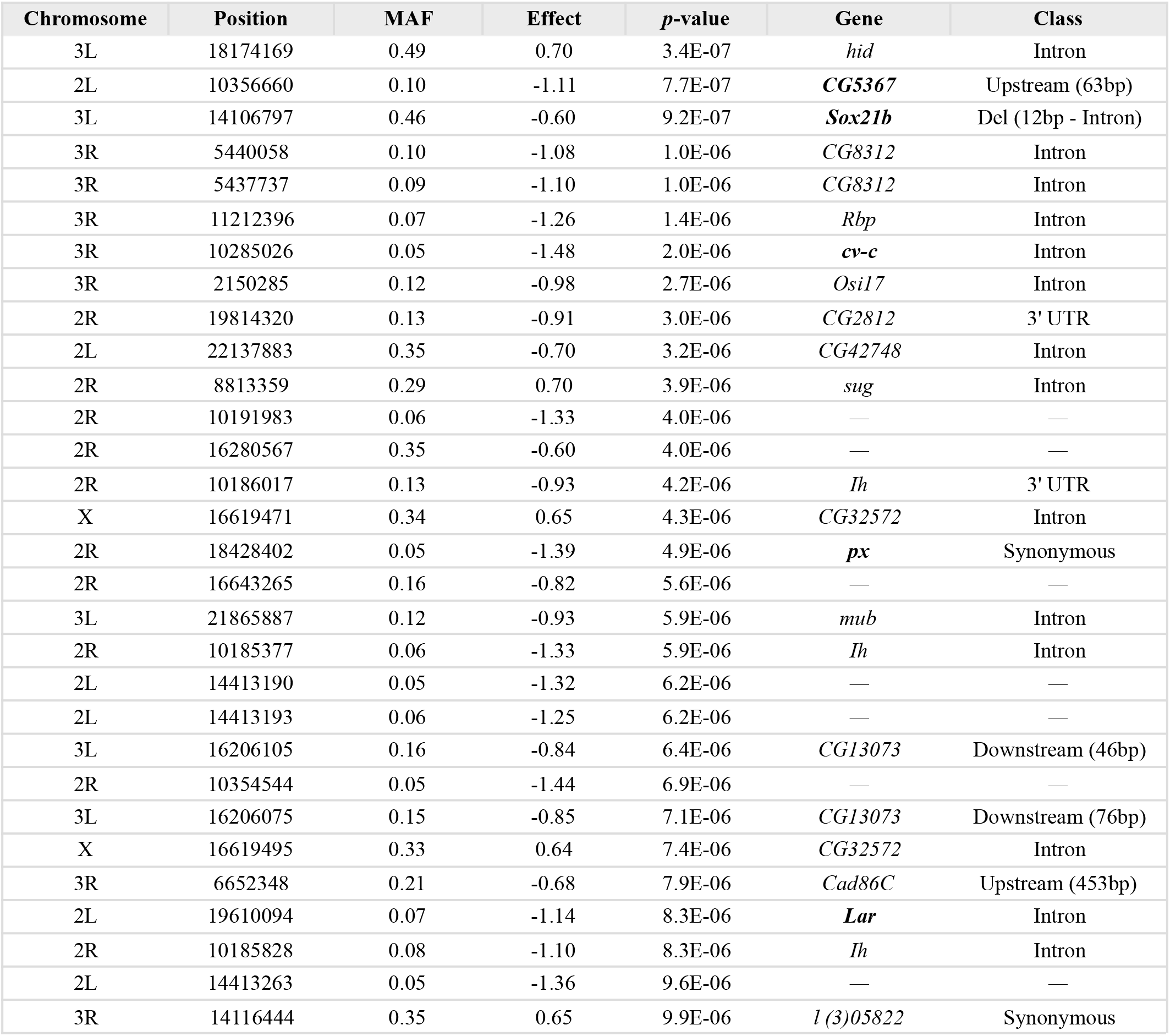
Variants significantly associated with the offspring index (*p* < 10^-5^). Chromosome coordinates represented in dm5 assembly coordinates. MAF = minor allele frequency, Del = deletion, and numbers in parentheses represent the number of basepairs to the closest gene. Genes in bold were previously identified to contain variants associated with age-specific fecundity^4^. Effect of variant on offspring index is unitless, as offspring index is a principal component score, derived from a PCA on scaled features.

### Offspring phenotype differences among lines are stable under different parental densities

Since we measured the variation in offspring index under a specific parental density during egg-laying, we decided to assess whether differences among DGRP lines in offspring phenotypes would persist under different parental densities. We chose six lines from our screen that were representative of negative offspring index (many low-weight offspring), intermediate offspring index, and positive offspring index (few high-weight offspring) to assay for offspring phenotypes at different densities of parents during egg-laying (Fig. 2). We found that parental density was a significant predictor of offspring number (*χ*^2^ test; females: *p* = 1.8×10^-7^, males: *p* = 4.7×10^-6^) and offspring weight (*χ*^2^ test; females: *p* = 2.2×10^-12^, males: *p* = 3.1×10^-12^). As expected, increasing parental density increased offspring number and decreased offspring weight, though the effect of increasing parental density increased sublinearly for most lines. After including density as a predictor, we saw that the DGRP line still had a significant impact on offspring number (*χ*^2^ test; females: *p* = 2.6×10^-5^, males: *p* = 1.8×10^-5^) and offspring weight (*χ*^2^ test; females: *p* = 9.5×10^-8^, males: *p* = 2.7×10^-7^). DGRP lines with positive index (RAL 812: +3.6; RAL 894: +3.8) maintained a low offspring number and high offspring weight under different densities. RAL 237, a DGRP line with a negative index (−2.9), had consistently high offspring numbers and low offspring weights. Surprisingly, RAL 176, a DGRP line with a strongly negative index (−3.2) yielded offspring with weights and counts similar to RAL 49, a line with an index close to zero (−0.04). We did not detect significant line-by-density interactions for any phenotype (*χ^2^* test; female number: *p* = 0.66, female weight: *p* = 0.86, male number: *p* = 0.54, male weight: *p* = 0.57).

**Figure 2.**
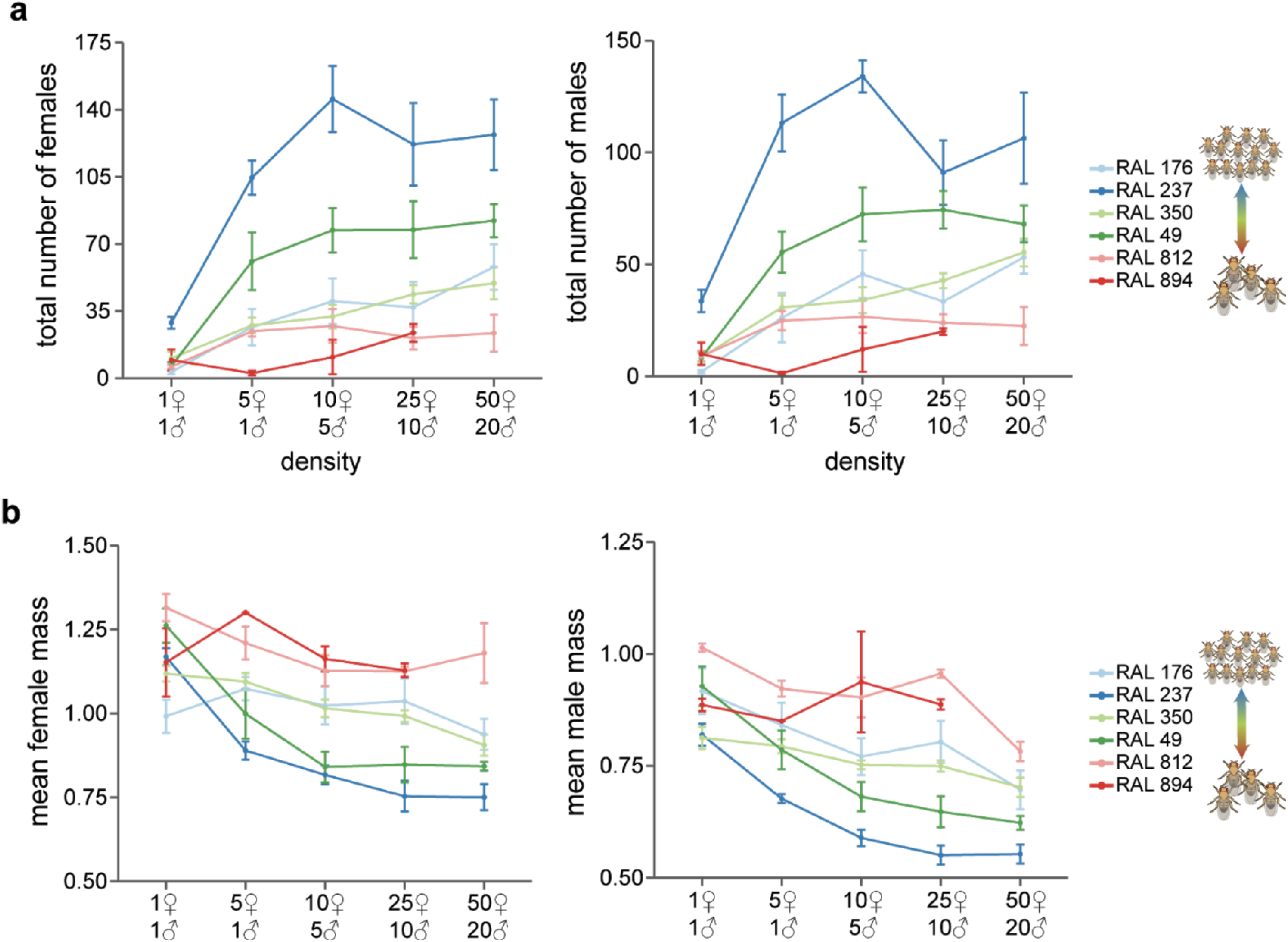
Independent effects of parental density and line on offspring number and weight. **a)** Relationship between number of offspring and the density of parents. Each point represents the mean phenotype for a line at a particular density and errors bars show +/− 1 s.e. (*n* = 2-7). **b)** Relationship between mean offspring weight and density of parents.

### Functional validation of associated variants

We chose six candidate genes to validate for involvement with our fecundity phenotype - *hid, Sox21b, Rbp, CG8312, mub,* and *Ih*. Genes for validation were chosen based on having a highly associated variant and availability of mutant lines. We used mutant lines from the Exelixis gene disruption panel, which contain *piggyBac* inserts in the genes of interest, to validate our candidate genes (Fig. 3). For four of the six candidate genes (*hid, Sox21b, CG8312,* and *mub),* we found that the available mutations severely impacted pupal and adult viability, to the point where we were unable to generate a stable homozygous line to use in our validation experiments (Table 2). With the remaining genes, *Rbp* and *Ih*, we found significant, opposite effects on the offspring index, with the *Ih* insertion strongly decreasing the offspring index, while the *Rbp* insertion slightly increased it (Fig. 3a). Examining component phenotypes (Fig. 3b), we saw that the *Ih* insertion significantly increased the number of offspring and decreased mean offspring weight, for offspring of both sexes. Disrupting *Rbp* did not have a statistically significant impact on the number of offspring, but there was a modest, but statistically significant, increase in offspring weight (13% for males and 21% for females). Both *Ih* and *Rbp* play a role in nervous system function. *Ih* encodes a voltage-gated potassium channel, and *Ih* mutants show defects in locomotion, proboscis extension, circadian rhythm, and lifespan^29,30^. *Rbp* encodes a protein involved in the organization of the presynaptic active zone and is instrumental in proper vesicle release^31^ - mutations in *Rbp* can result in neurological and locomotor defects, and in some cases, lethality.

**Figure 3.**
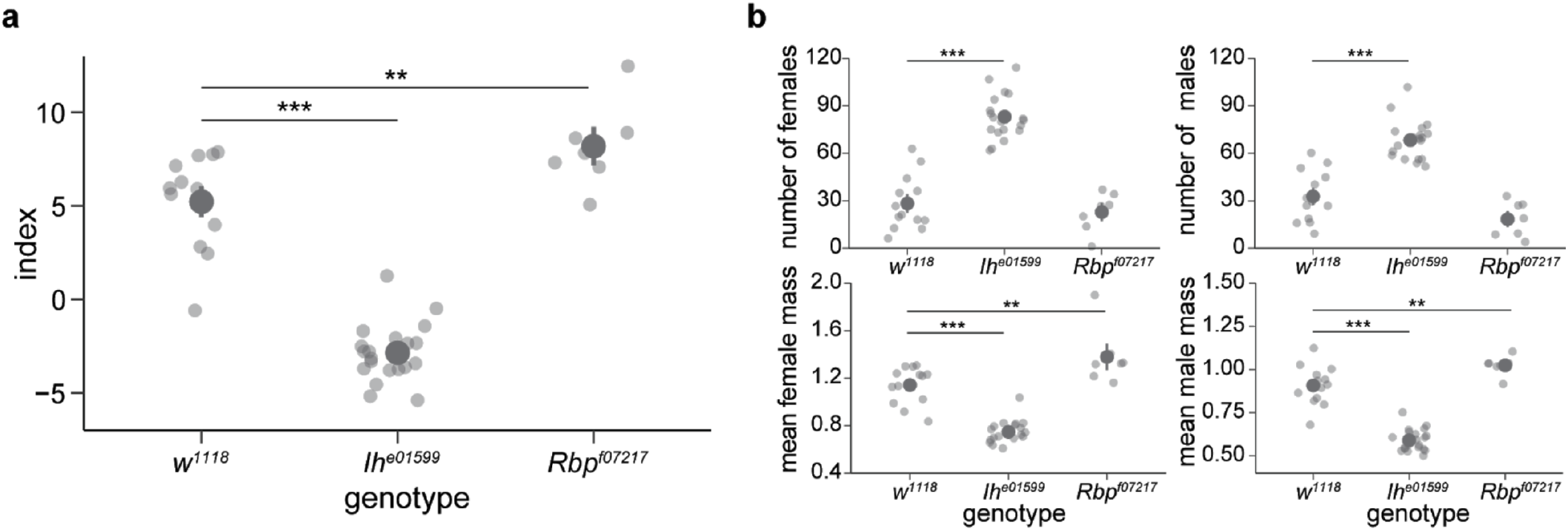
Candidate gene validation using *PBac{RB}Ih^e01599^* (n=10) and PBae{WH}*Rbp/^07217^* (n=7), compared to their genetic background control, *w^1118^*. **a)** Offspring index by genotype. **b)** Impact of gene disruption on individual phenotypes. In both panels dark gray points and bars show the mean +/− 1 s.e.; lighter gray points are replicates. Significance levels: * *p* < 0.05, ** *p* < 0.01, *** *p* < 0.001. *P*-values were calculated using Dunnett’s test with a family-wise confidence level of 95%.

**Table 2.**
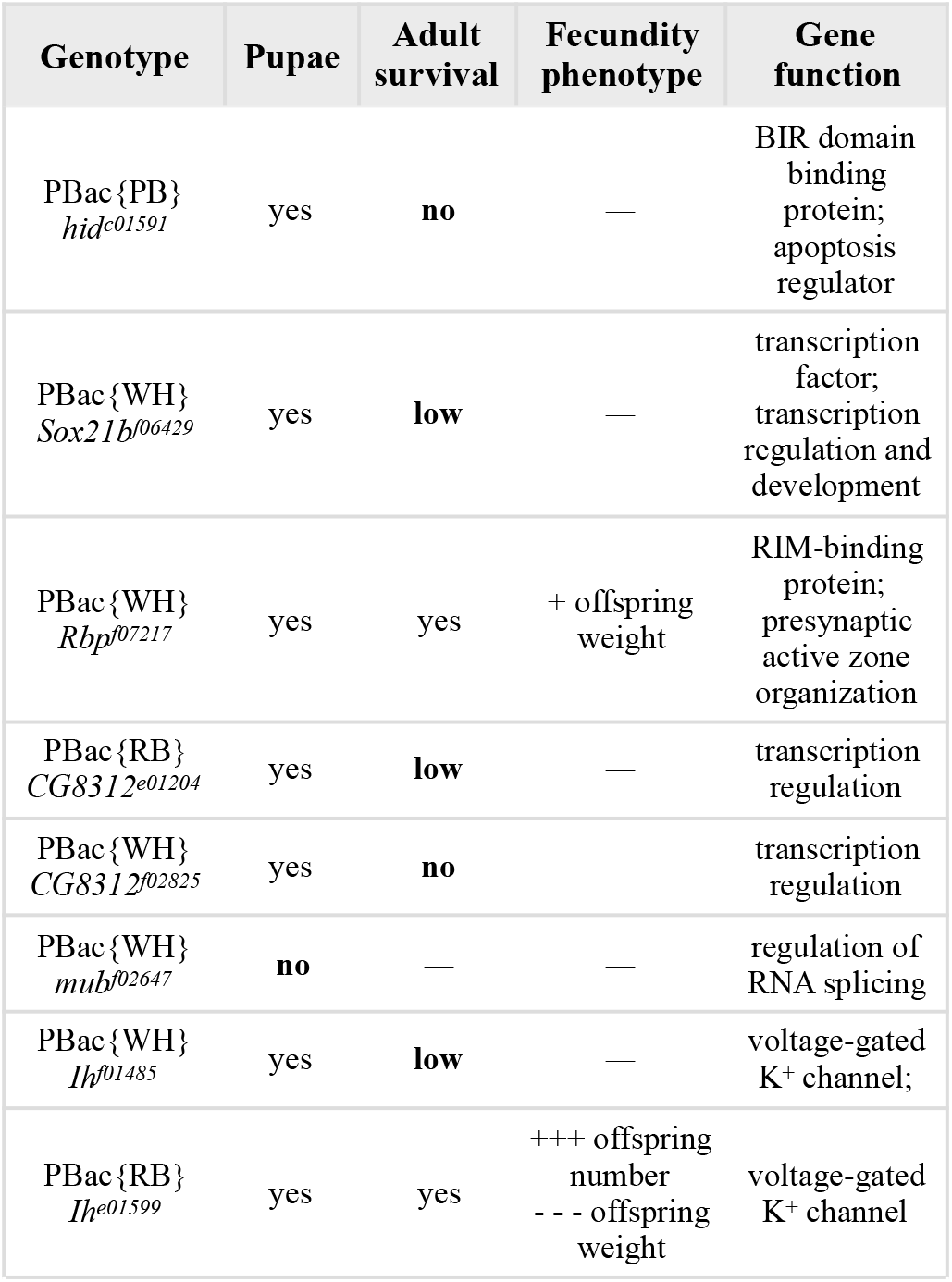
Homozygous mutant genotypes used in candidate gene validation and their fecundity phenotypes and gene functions. Gene function information was retrieved from FlyBase^33^. Number of + or − is a qualitative representation of how much the measured phenotype increased or decreased as compared to the control genetic background line (only statistically significant differences are shown).

For genes where we tested multiple mutant lines, we qualitatively compared the effect of insertion site on the fecundity phenotypes. We found that for *Ih, Ih^e01599^* was a viable line with increased offspring count, while *Ih^01485^* had impaired adult viability (Table 2). While both insertions are in introns, *Ih* transcription has been shown to be disrupted in *Ih^f01485^*, but not in *Ih^e01599^* ^29^. The effect of the insertion on the final protein function of *Ih^e01599^* is unknown, but RT-PCR shows that intronic insertion in *Ih^f01485^* results in a null mutation^32^.

### Correlations among candidate gene expression and traits

We examined whether the expression levels of candidate genes used in the validation were correlated in order to identify suites of co-regulated genes. We examined expression correlations among five of the six candidate genes (*CG8312* did not have available expression data). Using female expression data, we identified that *Ih* expression was positively correlated with *hid* expression and *Rbp* expression, following a multiple test correction (*Ih-hid: r* = 0.25,*p =* 2.5E-3; *Ih-Rbp: r* = 0.24,*p =* 3.1E-3) (Fig. 4a). Using the male expression data and a multiple test correction, we found that *mub* was positively correlated with *hid, Rbp,* and *Ih* and negatively correlated with *Sox21b* (Fig. 4b). We wanted to see whether the number of significant correlations we observed was higher than expected by chance. To this end, we sampled five random genes from the expressed data and calculated the number of significant correlations post-correction. We found that for both male and female expression, we have probably not enriched for a highly-correlated cluster of genes within our validation set (Fig. 4c-d). Using the same approach on a larger set of 17 candidate genes, we also did not find enrichment for gene expression correlations (Supp. Fig. 2).

**Figure 4.**
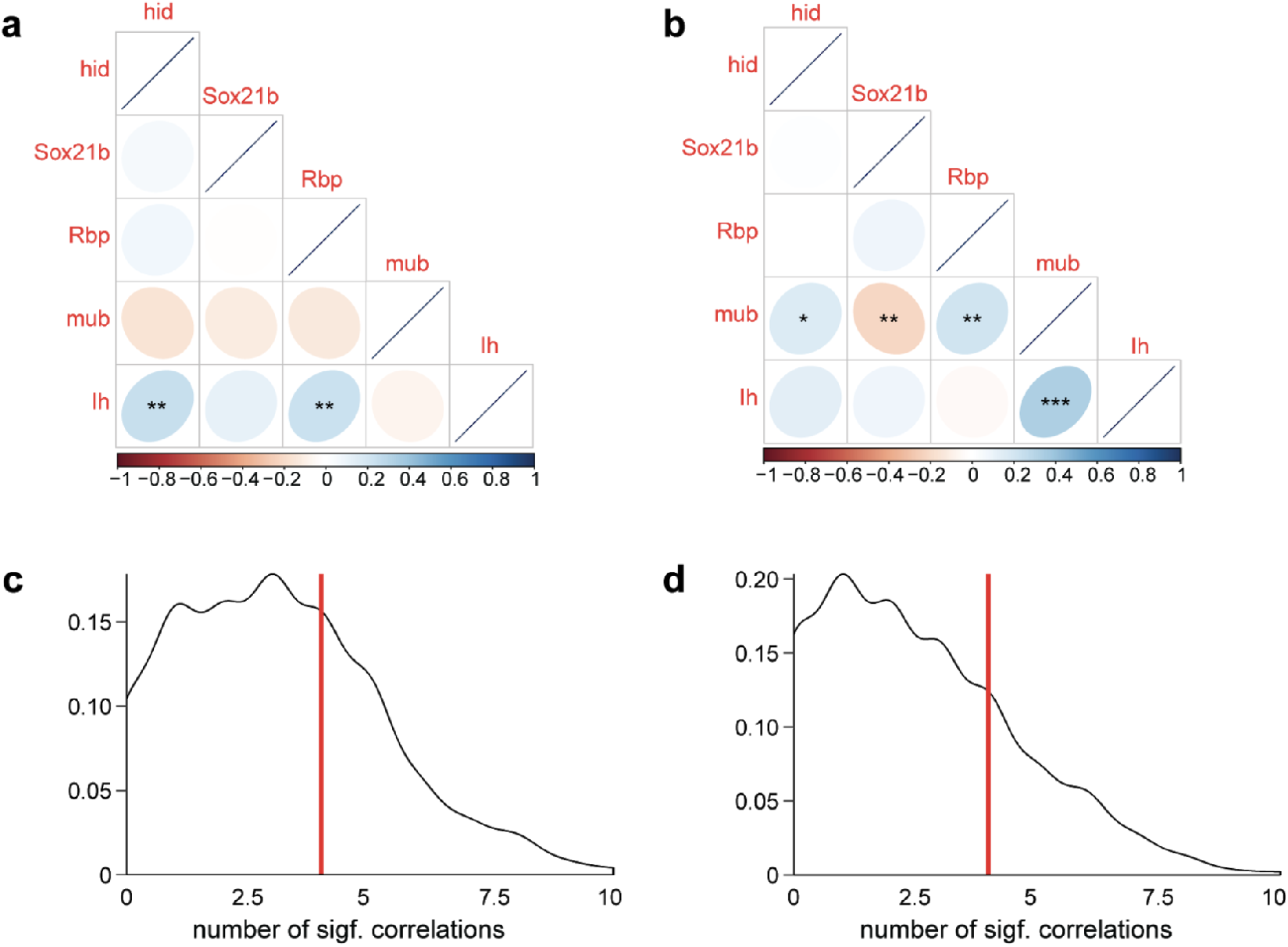
Correlation across DGRP lines of expression of candidate genes used in the validation experiment. **a)** Female expression data **b)** Male expression data. **c)** Kernel density plots of the number of significant correlations of expression of 5 randomly chosen genes (1000 samples) from female expression data. **d)** As in (c) for male expression data. Red line shows our observed number of significant correlations. Significance levels: * *q* < 0.05, ** *q* < 0.01, *** *q* < 0.001. *P*-values were transformed into *q*-values using the Benjamini-Hochberg method to correct for multiple tests. Here and below, *q*-values refer to the adjusted *p*-values that result from an FDR approach to multiple testing correction.

We also examined whether expression of the candidate genes was significantly correlated to the phenotypes measured in this study. We did not find evidence of strong correlations between the gene expression of our candidate genes and the phenotypes measured (Supp. Fig. 3).

Next we examined correlations between phenotypes measured in this study and those measured in other DGRP studies that might pertain to fecundity. We looked at the following phenotypes: starvation resistance^26^, chill coma recovery time^26^, food intake^34^, fecundity and body size^4^, nutritional indices and weight^25^, and developmental time and egg-to-adult viability under different densities^15^. Using a false discovery rate correction with all possible comparisons, we found that offspring weight measured in our study was significantly correlated with body size and mean weight measurements made in previous DGRP studies (Table 3). There was no correlation found between our measurements of total progeny number and fecundity measurements (*p* > 0.1 for all comparisons). In addition there was no correlation between progeny weight and food intake (*p* > 0.1 for all comparisons). We found nominally significant positive correlations between female starvation resistance and female (*r =* 0.18, *p =* 0.028) and male (*r =* 0.17, *p =* 0.046) weight measured in our study; these correlations were not significant after correcting for multiple tests. In addition, we found that male and female development time under a high larval density treatment (measured in 31 DGRP lines) was negatively correlated with the number of offspring, but positively correlated with offspring weight. Egg-to-adult viability under high larval density treatment was positively correlated with offspring number and negatively correlated with offspring weight. Though we observed a trend in the relationships between development time, viability and our phenotypes, only a few correlations were nominally significant, and none remained significant after the multiple testing correction.

**Table 3.** Correlations of phenotypes measured in our study (measured) with traits measured in previous DGRP studies (comparison). Only correlations that remained significant post-multiple testing correction are presented. Symbols are as follows: ‘-y’ denotes a low yeast diet, ‘+y’ denotes a high yeast diet, ‘+’ denotes a high glucose diet, ‘-’ denotes a low glucose diet, and ‘cb’ = overall effect when the data from the glucose diets are combined. *P*-values were transformed into *q*-values using the Benjamini-Hochberg method to correct for multiple tests across all comparisons.

| Measured | Comparison | $r$ | $q$-value | Ref |
| --- | --- | --- | --- | --- |
| mean weight (♂) | body size (-y) | 0.26 | 4.6E-02 | [4] |
| mean weight (♂) | body size (+y) | 0.28 | 2.9E-02 | [4] |
| mean weight (♀) | mean weight (♂-) | 0.41 | 3.5E-04 | [25] |
| mean weight (♂) | mean weight (♂-) | 0.39 | 6.9E-04 | [25] |
| total number (♂) | mean weight (♂+) | -0.30 | 2.9E-02 | [25] |
| mean weight (♀) | mean weight (♂+) | 0.44 | 1.4E-04 | [25] |
| mean weight (♂) | mean weight (♂+) | 0.41 | 3.5E-04 | [25] |
| mean weight (♀) | mean weight (♂<sup>cb</sup>) | 0.46 | 6.4E-05 | [25] |
| mean weight (♂) | mean weight (♂<sup>cb</sup>) | 0.43 | 1.4E-04 | [25] |

## Discussion

We investigated whether there was a genetic basis for adult offspring number and weight under space and resource limitation. We found that DGRP lines varied in the number of offspring produced and their mean weight, with number negatively correlated with weight. Using a combined ‘offspring index’ derived from the first principal component of our offspring phenotypes, we identified variants associated with variation in offspring index (i.e., variation from many low-weight offspring to few high-weight offspring). We examined the effects of gene disruption on six candidate genes and found that for all tested insertion alleles of *hid, Sox21b, CG8312,* and *mub,* as well as one allele of *Ih*, gene disruption caused phenotypes ranging from pupal lethality to low adult survival. Disruption of *Rbp* caused a small increase in offspring weight and an insertion allele of *Ih* caused a large increase in offspring number coupled with a decrease in offspring weight. While we did find significant correlations in gene expression between the candidate genes, the number of significant correlations did not exceed what we would expect from randomly selected genes. When comparing our measured phenotypes to life-history phenotypes measured in other DGRP studies, we found consistency in our body weight measurements and other measurements of weight and body size, but surprisingly, we did not find a relationship between offspring number and previous measures of fecundity.

We found that, similar to fecundity as measured by number of eggs laid^4^, there is a polygenic basis to offspring number and weight. While several of the candidate genes we found were previously associated with fecundity^4^, most were not. Among our candidate genes, we were not able to find significant enrichment of genes involved in particular biological processes or molecular functions, though given our limited sample size of genes, only a very strong enrichment could have been significant. While only two of the six of the candidate genes tested in our validation experiments are annotated as having roles in developmental processes, we found that for most genes tested, mutation disrupted the survival of either the larval, pupal or adult stages severely enough to prevent us from even measuring offspring number and weight.

The omnigenic model of genetic architecture^35,36^ could explain the functionally varied suite of genes associated with offspring number and weight phenotypes. One of the predictions of the omnigenic model is that essentially all genes contribute to a trait through their expression at some appropriate developmental point or in a particular tissue. This leads to many loci in diverse genes being weakly associated with the trait of interest. The omnigenic model also predicts that there should be a network of genes whose action is essential and that have correspondingly higher effect sizes. Interestingly, even though we observed that disruptions to our candidate genes led to strong effects, we did not find an enrichment for correlated expression among them, i.e., they did not appear to be part of a transcriptional network. We were also unable to find enrichment for particular biological pathways or molecular processes among our candidate genes. Since we were unable to identify a network uniting our candidate genes, our results are not consistent with an omnigenic model, but perhaps reflect what was proposed in Fisher’s “infinitesimal model”^37^, in which a quantitative trait is made up of infinitely small contributions of infinitely many genes.

We found that insertion mutations in *Rbp* and *Ih* caused opposite phenotypes. Mutations in *Ih* increased the offspring number and, presumably due to the constraints of the vial, decreased offspring weight. *Ih* had not been previously implicated in fecundity phenotypes, as measured by egg laying. While that does not preclude that *Ih* mutants may lay more eggs, a higher egg-to-adult viability for *Ih* mutants could also lead to the increase in offspring number. Disruption of *Rbp* only increased weight without significantly affecting offspring number. A small effect on offspring number could have been obscured by limited sample size, but our results still indicate that the weight increase was more prominent than the decrease in offspring number. A decoupling of weight from offspring number shows that there is an independent axis where offspring weight can increase even though the level of larval competition and other density effects remain the same. This notion is supported by the second principal component of our dataset which is positively loaded for both weight and offspring number, indicating that a trade-off between the two is not mandatory (Supp. Table 2).

We observed that even within the same gene, different disruptions can lead to different phenotypes. Both the *Ih^e01599^* and *Ih-f^01485^* alleles are intronic insertions that affect most transcripts^29,32^, but *Ih^f01485^* shows impaired viability, while *Ih^e01599^* shows an increase in offspring number. The *Ih^f01485^* allele was reported to eliminate expression of all *Ih* transcripts^31^. In contrast, the *Ih^e01599^* allele was reported as having wild-type levels of expression^29^. Based on its insertion position, the *Ih^e01599^* allele would affect 8 of the 11 *Ih* transcripts. The phenotype difference we observed between these alleles could depend on specific transcripts. These results highlight a general caveat about using mutant lines to make conclusions about the exact role of the gene in determining the phenotype, rather than a more general conclusion about whether or not the gene plays a role at all.

When we collected offspring from different parents at different densities, we found a sub-linear increase in the number of progeny, consistent with the effects of resource limitation. We also found that offspring phenotypes were strongly determined by DGRP line at all densities. To first approximation, DGRP lines that produce fewer larger offspring continued to produce fewer larger offspring regardless of the number of parents allowed to oviposit in the vial. There appear to be parallels between the variation in offspring phenotypes among DGRP lines and *r* and *K-*selection^38^. In an ecological context, an *r-*selected species is one that produces many offspring with low chances of survival. A *K-*selected species invests in few high-quality offspring in order to be able to compete in more crowded environments with limited resources. If the *r/K* framework plays out as a plastic trait within a genotype, we might expect lines to converge on smaller offspring in conditions of higher crowding. We observed no such convergence, nor indeed any significant line-by-density effects. The genetically determined variation in offspring phenotypes among our lines seems to reflect a within-species continuum between *r* and *K-*selected types, perhaps consistent with a Pareto frontier of equivalently fit phenotypic combinations. Given that we are studying this trait in a collection of inbred lines meant to capture the genetic variation of an outbred population, it is also possible that outbred *D. melanogaster* populations exhibit less variation on this continuum, while the inbred lines lock in diverse phenotypes based on their varied genetic compositions.

Comparing our phenotypes to those measured in other DGRP studies, we found correspondence between our weight measurements and body weight/size measurements from other studies, affirming that DGRP line phenotypes can remain consistent across different study environments. We did not, however, see a relationship between the number of offspring and prior fecundity measures. This is unexpected, as one would expect that the number of eggs laid should correlate with the number of offspring. This may be indicative of substantial line-to-line variation in mortality post-egg-laying but pre-eclosion. Another possible explanation for the lack of correlation could be due fecundity being previously assayed with individual females^4^, whereas we housed females in groups of 10 for our assay. The number of eggs laid per female was shown to decrease in more crowded conditions^13,39^, and with the presence of differential genotype effects^39^, this could lead to the lack of correlation observed. We also observed that, within 31 DGRP lines, more offspring is correlated with a lower developmental time and higher egg-to-adult viability under a high density treatment. Though these correlations were not significant after a multiple testing correction, it does indicate that lines producing more offspring may have adapted to do well under high-density conditions^15,19^.

Overall, our results point to a polygenic basis for offspring number and weight. Validation of six candidate genes implicates diverse biological processes in controlling adult stage viability. Combining our results with results from studies on other *Drosophila* life-history traits, we find support for the idea that traits closely related to fitness (i.e., offspring number) may be influenced by a large set of genes, perhaps ultimately encompassing the vast majority of functional genes.

## Methods

Data and analysis code are available at *https://zenodo.org/record/3932230* and *http://lab.debiyort.org/genetic-basis-of-off-spring-number/*.

### Drosophila stocks and husbandry

We analyzed 143 lines from the *Drosophila* Genetic Reference Panel (DGRP^26^). All stocks were maintained in incubators at 23°C, 12L:12D cycle and reared on a yeast, cornmeal, and dextrose media (23g yeast/L, 30g cornmeal/L, 110g dextrose/L, 6.4g agar/L, and 0.12% Tegosept). Experiments were carried out in polystyrene narrow culture vials (25 x 95 mm, #32-109, Genesee Scientific). Mutant lines for validation were obtained from the Exelixis collection at Harvard Medical School (Boston, MA, U.S.A.)^40^. The *w^1118^* (#6326) genetic background was obtained from the Bloomington Drosophila Stock Center (Bloomington, IN, U.S.A.) as a control for the candidate gene validation mutant lines.

### Phenotypic measurements of DGRP lines

Bottles were seeded with 15 females and 15 males in 12L: 12D to generate the parent flies that would go on to lay eggs for the experiment. Ten females and five males (2-5 days old) from the parent flies were placed in each of the three vials (along with ~30 grains of dry yeast) and left to oviposit for 2 days at 23°C, 12L:12D. The parent flies were removed and the vials were kept at 23°C, 12L:12D until progeny began to eclose. From the start of eclosion, all of the vials were examined every day over the course of 10 days. The number of females and males for each vial was recorded, as well as the total wet weight (to 0.1 mg accuracy) of the females and the total combined wet weight of the females and males. Male weight was calculated by subtracting the female weight from the total. Ten days was chosen to measure as many offspring as possible without measuring any flies from the subsequent generation. 143 lines were tested in two batches. 134 lines were tested in the first batch. The second batch included the 35 lines that did not have all three vial replicates completed in the first batch, and 9 lines that were not tested at all in the first batch.

### Genome-wide association mapping for offspring index

The four phenotypes measured in this screen were the total number of females (males) eclosed and the average weight of a female (male) flies. Since lines were assayed in two batches, the 35 lines that were tested in both batches were used to check for a batch effect. The batch effect for each of the four phenotypes was corrected by applying an offset (difference of mean phenotype between the first and second batch) calculated from the overlapping set of lines: (number of ♀ = −17, number of ♂ = −22, mean ♀ weight = 0.13, mean ♂ weight = 0.12).

For offspring total counts, a random intercept linear model was used to calculate the random effect of each DGRP line:

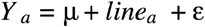

where *Y_a_* is the phenotype measure for line *a*, linea is the random effect of line *a*, and is the error term. For mean offspring weights, a random intercept generalized linear model was used (model formula as above), assuming a gamma distribution of mean weights and a logarithm link function. The LME4 package (v1.1-21) in R (v3.5.3) was used for modeling. To estimate heritability for each phenotype, we used the R package brms (v2.8.0) and our models to estimate the fraction of variance explained by line out of all sources of variance, as well as the uncertainty in the estimate from the 95% highest posterior density interval.

Since we were interested in a single metric to summarize the number of offspring and their average weight, we used *prcomp* with scaling in R’s factoextra package (v1.0.5) to generate the principal components of the dataset. The first principal component explained 71% of the variance of the dataset, so we chose to use the value of the rotated data (line phenotype values multiplied by the rotation values/loadings of the first principal component) as our summary phenotype, which we called the offspring index (Supp. Table 2).

We used the DGRP2 webtool^41^ to perform a mapping of variants associated with the offspring index. The webtool controls for inversions and *Wolbachia* infection status prior to mapping. We chose a significance threshold of *p* < 10^-5^ to identify variants for further consideration. A *χ*^2^ test test was run to compare the observed chromosomal distribution of variants to the expected distribution given the proportion of all segregating variants on each chromosome in the DGRP dataset^41^.

### Parental density analysis

Six DGRP lines were chosen from the overall screen based on their offspring index values: two lines with an extreme negative index (RAL 176: −3.2, RAL 327: −2.9), two lines with an index around zero (RAL 49: −0.04, RAL 350: −0.22), and two lines with a highly positive index (RAL 812: +3.6, RAL 894: +3.8). Five density conditions were used: 1♀,1♂, 5♀,1♂, 10♀,5♂, 25♀,10♂, 50♀,20♂. Lines were reared in bottles at 23°C, in incubators on a 12L:12D light cycle, and a minimum of 5 (maximum of 10) replicates per line-density combination were set up, with the exception of RAL 894 at the highest density, where no replicates were set up due to an insufficient number of flies. Parent flies were 3-5 days old at the time of experiment set-up and the egg-laying conditions were the same as the DGRP phenotype screen. Given the delayed eclosion of offspring in the high density treatments, an extended window of 25 days was used to evaluate offspring phenotypes. Daily records after the 12^th^ day were monitored for increases in offspring number that could be indicative of a large number offspring from the next generation, but as flies were removed daily from the vials, likely even before mating, a significant influence of next-generation offspring on the counts was deemed unlikely.

To estimate the impact of DGRP line and density on offspring counts, we used a generalized linear model, assuming a negative binomial distribution of the response along with a logarithm link function. A negative binomial model was used because of the large spread and right-skew of the offspring count distribution that made a regular linear model a poor fit. For offspring weight, a generalized linear model with a gamma response distribution and logarithm link function was used. The formula for both models was as follows:

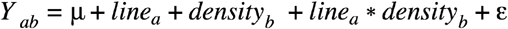

where *Y_ab_* is the phenotype measure for a particular line-density combination, *linea* is the fixed effect of line *a*, *densityb* is the fixed effect of a density treatment *b*, *linea*densityb* is the interaction term, and is the error term. DGRP line and density were treated as ordered factors in the model. DGRP lines were treated as ordered factors since the lines for this experiment were chosen to span the range of offspring index values. The lines were coded from 1-6: line 1 with the most negative offspring index and line 6 with the most positive offspring index. As above, the LME4 package was used, in addition to the MASS package (v7.3-51.1) for the negative binomial model. To evaluate the significance of a predictor, a likelihood ratio *χ*^2^ test using the *anova* function in R’s STATS (v3.5.3) package was used to compare models with and without the predictor.

### Validation of candidate genes

Mutant lines for validation of candidate genes were obtained from the Exelixis collection^24^ for six genes containing variants associated with the offspring index at a *p* < 1E-5 threshold. The mutant lines were as follows: PBac{PB}*hid^c01591^*, PBac{WH} *Sox21b^f06429^,* PBac{WH}*Rbp^f07217^*, PBac{RB}*CG8312^e01204^*, PBac{WH}*CG8312^f02825^,* PBac{WH}*mub^f02647^*, PBac{WH}*Ih-^f01485^,* and PBac{RB}*Ih^e01599^* (#17970 BDSC). All lines were made homozygous for the insertion prior to testing. The genetic background for this gene disruption panel was *w^1118^*. To generate the parental flies for each mutant line and the control, ~30 females and 10 males were placed in bottles at 22°C, 12L:12D to lay for 7-10 days to generate the experimental flies. 10 females and 5 males from the experimental flies were put into a single vial with ~30 yeast grains (ten replicates per mutant line) and allowed to lay for 2 days at 22°C, 12L:12D. The parental flies were removed and the progeny were phenotyped the same way as for the DGRP phenotype screen. The validations were done in two batches staggered by one week. There was no significant batch effect, so replicates were combined across batches. Mutant lines were compared to the *w^1118^* genetic background control using Dunnett’s test with a family-wise confidence level of 95%. Offspring index for the mutant lines was calculated using the principal component loadings calculated from the DGRP data.

### Gene expression correlations and correlations with other traits

Gene expression data was obtained from Huang *et al.* (2015)^42^. We calculated the Pearson correlation between the expression of candidate genes tested in the validation experiment, as well as the correlation between the expression of those genes and the phenotypes measured in our screen. We also correlated the traits measured in this study against similar or potentially related traits measured in other DGRP studies^4,25,26,34^. All analysis was done in R (v3.5.3).

## Acknowledgements

We thank Rob Unckless and Alan O. Bergland for sharing raw data on DGRP phenotypes. B.L.d.B. was supported by a Sloan Research Fellowship, a Klingenstein-Simons Fellowship Award, a Smith Family Odyssey Award, and National Science Foundation grant no. IOS-1557913. J.A. was supported by Harvard’s Quantitative Biology Initiative and National Science Foundation PoLS-HF-SRN (NSF-1806818).

## Conflicts of Interest

The authors declare no competing interests.

## Supplementary materials

**Supp. Table 1.** Heritability estimates for the means of the four phenotypes. HPD = highest posterior density.

| Phenotype | Heritability Estimate | 95% HPD Interval |
| --- | --- | --- |
| Mean ♀ Weight | 0.64 | 0.55 - 0.72 |
| Mean ♂ Weight | 0.73 | 0.65 - 0.80 |
| Number of ♀ | 0.47 | 0.37 - 0.57 |
| Number of ♂ | 0.48 | 0.39 - 0.58 |

**Supp. Table 2.** Variance proportion explained by each principal component and their loadings.

|  | PC1 | PC2 | PC3 | PC4 |
| --- | --- | --- | --- | --- |
| **St. dev** | 1.69 | 0.97 | 0.39 | 0.28 |
| **Variance prop.** | 0.71 | 0.23 | 0.037 | 0.020 |
| **Number of ♀** | -0.51 | 0.50 | -0.19 | 0.68 |
| **Number of ♂** | -0.51 | 0.48 | 0.21 | -0.68 |
| **Mean ♀ Weight** | 0.50 | 0.49 | 0.69 | 0.19 |

**Supp. Figure 1.**
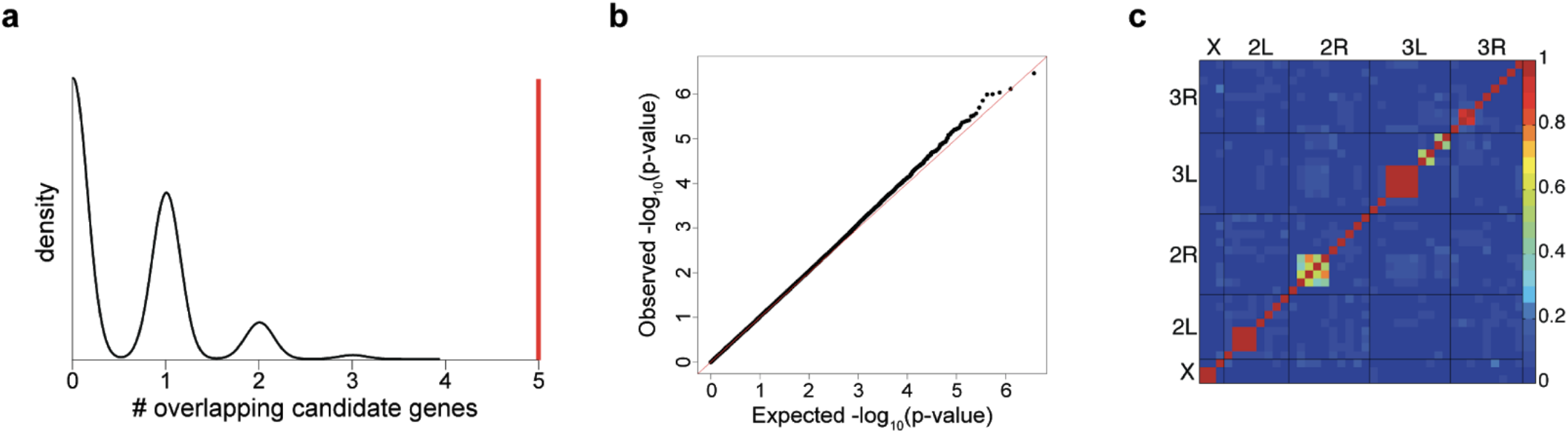
**a)** Bootstrap distribution (*n* = 1000) of the number of overlapping genes between our candidate genes and a random set of candidate genes (black kernel density estimate) vs. the observed number of overlapping genes (red line) between our candidate genes and 552 candidate genes previously associated with fecundity4. For each bootstrap sample, a random set of 552 candidate genes was chosen without replacement from a list of genes associated with DGRP variants (annotation based on FlyBase release 5.57). **b)** QQ plot comparing observed and expected *p-*values, with the red line showing a 1:1 relationship. **c)** Linkage disequilibrium heat map for all the variants identified (*p* < 10E^-4^).

**Supp. Figure 2.**
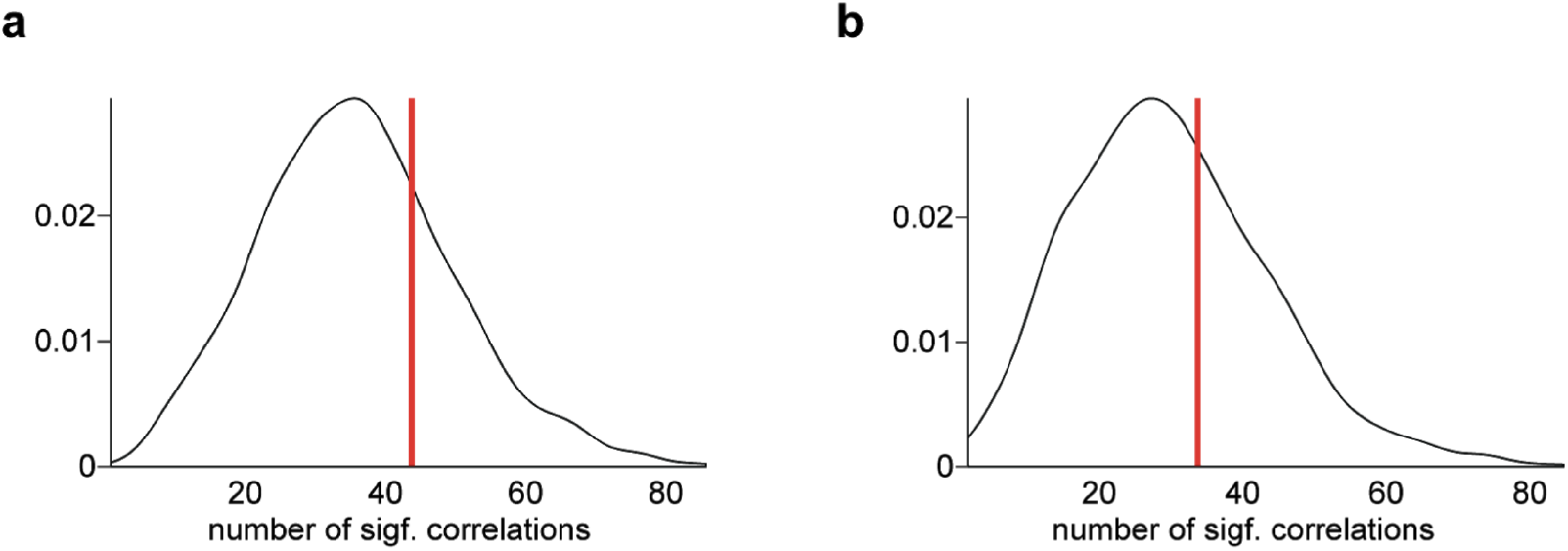
Kernel density plots of the number of significant correlations in expression among 17 randomly chosen genes. (1000 resamples were computed) from a) female expression data and b) male expression data. Red line shows our observed number of significant correlations among 17 candidate genes examined. *P*-values were transformed into *q*-values using the Benjamini-Hochberg method to correct for multiple tests prior to determining significance (FDR = 0.05). *Q*-values refer to the adjusted *p*-values that result from an FDR approach to multiple testing correction.

**Supp. Figure 3.**
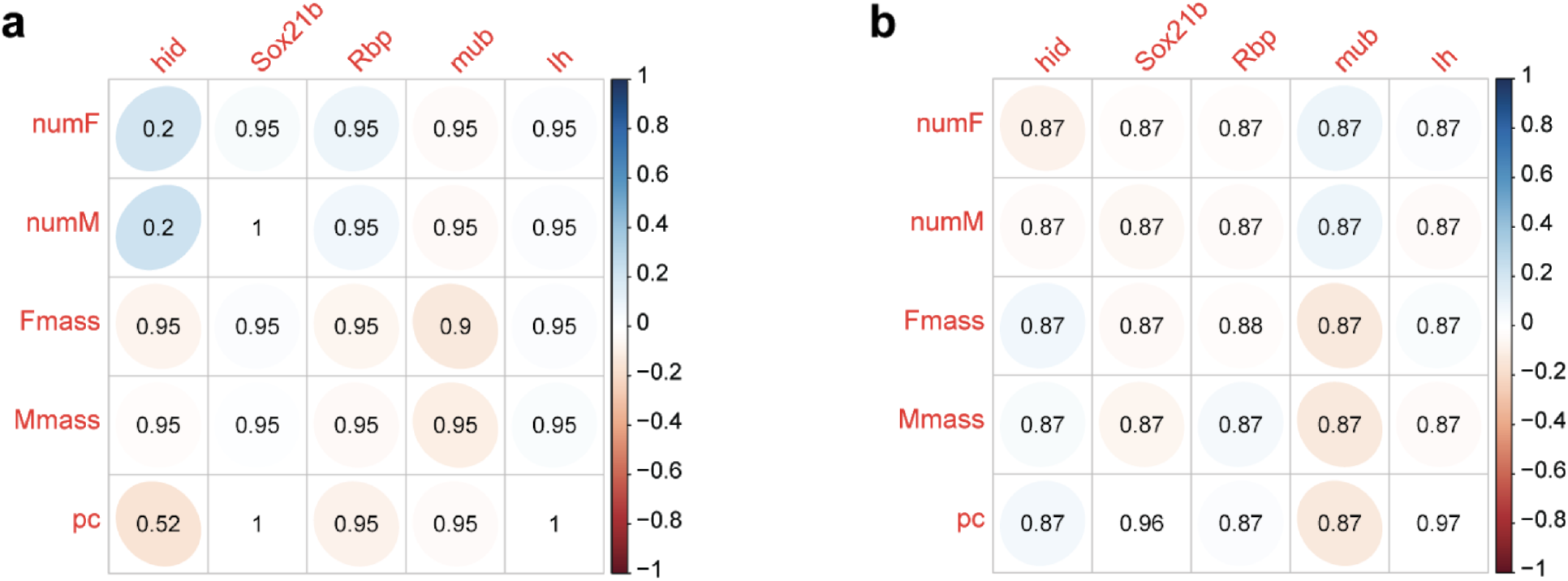
Correlation of expression of candidate genes to offspring phenotypes. **a)** DGRP female expression data **b)** DGRP male expression data. pc = offspring index. *P*-values were transformed into *q*-values using the Benjamini-Hochberg method to correct for multiple tests. *Q*-values refer to the adjusted *p*-values that result from an FDR approach to multiple testing correction.

## Notes

https://zenodo.org/record/3932230

http://lab.debivort.org/genetic-basis-of-offspring-number/

### Competing Interest Statement

The authors have declared no competing interest.

